# Time-series transcriptomics from cold, oxic subseafloor crustal fluids reveals a motile, mixotrophic microbial community

**DOI:** 10.1101/2020.08.03.235341

**Authors:** L.M. Seyler, E. Trembath-Reichert, B.J. Tully, J.A. Huber

## Abstract

The rock-hosted oceanic crustal aquifer is one of the largest habitable volumes on Earth, and it harbors a reservoir of microbial life that influences global-scale biogeochemical cycles. Here, we use time series metagenomic and metatranscriptomic data from a low-temperature, ridge flank environment that is representative of the majority of global hydrothermal fluid circulation in the ocean to reconstruct microbial metabolic potential, transcript abundance, and community dynamics. The data suggest that the microbial community in this subseafloor habitat is motile, chiefly heterotrophic or mixotrophic, and capable of using alternative electron acceptors such as nitrate and thiosulfate, in addition to oxygen. Anaerobic processes are most abundant in subseafloor horizons deepest in the aquifer, furthest from connectivity with the deep ocean, and there was little overlap in the active microbial populations between sampling horizons. Together, our results indicate the microbial community in the North Pond aquifer plays an important role in the oxidation of organic carbon within the crust, and is also metabolically flexible, with the ability to switch from autotrophy to heterotrophy, as well as function under low oxygen conditions. This work highlights the heterogeneity of microbial life in the subseafloor aquifer and provides new insights into biogeochemical cycling in ocean crust.

## Introduction

Seawater circulating through porous, basaltic oceanic crust constitutes the largest actively flowing aquifer system on Earth, and represents approximately 2% of the ocean fluid volume (∼27 million km^3^ of water; [1, 2, 3]). The convective circulation of seawater through mid-ocean ridges and ridge flanks has profound effects on crustal and ocean chemistry [4, 5]. Studies of warm, anoxic venting fluids from mid-ocean ridges and ridge flanks indicate a diverse and active microbial community in these crustal fluids (reviewed in [6]). However, much of the remaining crustal pore volume is composed of cold, oxygenated deep ocean water that enters and exits the basaltic crust through seafloor exposures [7, 8]. These crustal fluids are chemically similar to seawater, but slightly enriched in DIC and depleted in DOC and O_2_ [9, 10, 11, 12]. Microbial communities in the crustal fluids are distinct from those in the surrounding bottom water [9, 13], but the lifestyle and adaptive strategies of microbial communities and the biogeochemical processes they mediate in the oligotrophic, oxic subseafloor aquifer remain poorly constrained.

North Pond is an 8 km x 15 km sediment-filled basin located in 8 Mya volcanic crust on the western flank of the Mid-Atlantic Ridge, ∼4,450 m below the oligotrophic Sargasso Sea [14, 15, 16]. The overlying low-permeability sediments prevent seawater flow into the crustal aquifer in the porous and permeable basaltic crust below. In 2011, two circulation obviation retrofit kits (CORKs [17, 18]) were installed into drill holes U1382A and U1383C which penetrated oceanic crust at North Pond during IODP Expedition 336, thus enabling sampling and monitoring of the crustal aquifer [19]. The first crustal fluids were collected from the North Pond CORKs six months after installation in 2012, with return expeditions in 2014 and 2017. In addition, a battery-powered GeoMICROBE sled was deployed at each CORK for autonomous time series sampling from April 2012 to April 2014 [20]. 16S rRNA gene sequencing from 2012 showed that the North Pond fluid microbial community is distinct from background seawater and U1382A and U1383C communities are distinguishable from one another [9]. Metagenomic data from samples collected between 2012 and 2014 revealed large shifts in the dominant taxonomic groups, with a high degree of functional redundancy in the microbial community [13].

Natural abundance isotopic data from fluids sampled in 2012 and 2014 indicate early removal of dissolved organic carbon (DOC) sourced from the deep ocean, followed by the slower removal of older, more refractory components, with limited chemosynthetic production in the deepest basement fluids [10]. Experiments using fluids collected in 2012 incubated with ^13^C-labeled bicarbonate and acetate at 5°C and 25°C detected both autotrophic and heterotrophic activity at a higher level than in bottom seawater, [9], but low rates of metabolic activity were detected in these same fluids at 4°C using nanocalorimetry [21]. However, all of these samples were taken before it was clear the North Pond aquifer had chemically rebounded from the initial drilling disturbances, and time series geochemical data suggests that by 2017, the North Pond aquifer had recovered [12]. Therefore, fluids collected from North Pond in 2017 are likely the most representative of microbial communities in the cold, oxic aquifer.

Here, we present the first metatranscriptomic data from North Pond, including samples from 2012, 2014, and 2017, analyzed to reconstruct microbial metabolic potential, transcript abundance, and community dynamics in the cool, oxic marine crustal habitat. We compare newly sampled metagenomes from 2017 and metatranscriptomes across all three sampling years, with an emphasis on understanding the 2017 population as it represents the community furthest from drilling disturbances. We demonstrate the presence and activity of a motile microbial community with considerable metabolic flexibility, including the ability to carry out both autotrophy and heterotrophy under oxic and anoxic conditions, and present a conceptual model for the key microbially-mediated carbon, nitrogen, and sulfur cycling reactions occurring in the North Pond crustal fluids.

## Methods

### Sample Collection

Fluids were collected from the North Pond crustal aquifer (22°45′N and 46°05′W) in April 2012, April 2014, and October 2017 using ROV JASON and the Mobile Pumping System (MPS [20]) as described elsewhere [9, 13, 18, Trembath-Reichert et al. In Review]. Both U1382A and U1383C CORK installations were sampled via umbilicals that are accessible at the seafloor and terminate at different depths beneath the seafloor, as shown in Table 1 and Figure 5. CORK observatory U1382A contains a packer seal in the bottom of the casing that isolates the aquifer from the overlying sediment and one sampling depth horizon extending 90-210 meters below seafloor (mbsf). Observatory U1383C contains three sampling depth horizons below the sediment-crust interface separated by packer seals: Shallow (∼70-146 mbsf), Middle (146-200 mbsf), and Deep (∼200-332 mbsf). In all years, umbilical lines were flushed before sampling began and both temperature and oxygen were monitored throughout sampling [9, 12, 13]. To collect microbial biomass for -omics analyses in 2017, crustal fluid was pumped at a rate of ∼0.5 LPM for 80 min through a 0.22 µm, 47 mm GWSP filter (Millipore). These filters were fixed on the seafloor using RNAlater™ (Ambion) as described in [13, 20]. Bottom seawater was also collected at 4,397 m water depth using the MPS. Shipboard, all filters were placed in fresh RNALater™, incubated at 4 °C for 18 hours, and then stored at -80 °C until extraction.

**Table 1.** Samples analyzed with cell enumeration and oxygen data. *Depth in meters below surface, remaining depths are in meters below seafloor (mbsf).

| **Sample** | **Date Collected** | **Depth (m)** | **Cells per ml<sup>1</sup>** | **Oxygen (μM)<sup>2</sup>** |
| --- | --- | --- | --- | --- |
| U1382A | 25-Apr-2012 | 90-210 | $1.4 \times 10^4$ | 244 |
| U1382A | 5-Apr-2014 | | $2.8 \times 10^4$ | 233 |
| U1382A | 12-Oct-2017 | | $5.1 \times 10^3$ | 228 |
| U1383C Shallow | 30-Apr-2012 | 70-146 | $2.0 \times 10^4$ | 216 |
| U1383C Shallow | 2-Apr-2014 | | $1.1 \times 10^4$ | 202 |
| U1383C Shallow | 13-Oct-2017 | | $3.7 \times 10^3$ | 198 |
| U1383C Middle | 30-Apr-2012 | 146-200 | $2.2 \times 10^4$ | n.d. |
| U1383C Middle | 15-Oct-2017 | | $2.1 \times 10^3$ | 205 |
| U1383C Deep | 20-Apr-2012 | 200-332 | $2.1 \times 10^4$ | 213 |
| U1383C Deep | 31-Mar-2014 | | $2.8 \times 10^4$ | 187 |
| U1383C Deep | 11-Oct-2017 | | $3.7 \times 10^3$ | 173 |
| Bottom Water | 14-Oct-2017 | 4397* | $9.0 \times 10^3$ | 250 |

<sup>1</sup>Cell counts from [13] and Trembath-Reichert et al., In Review.
<sup>2</sup>Oxygen data from [12].

**Figure 1.**
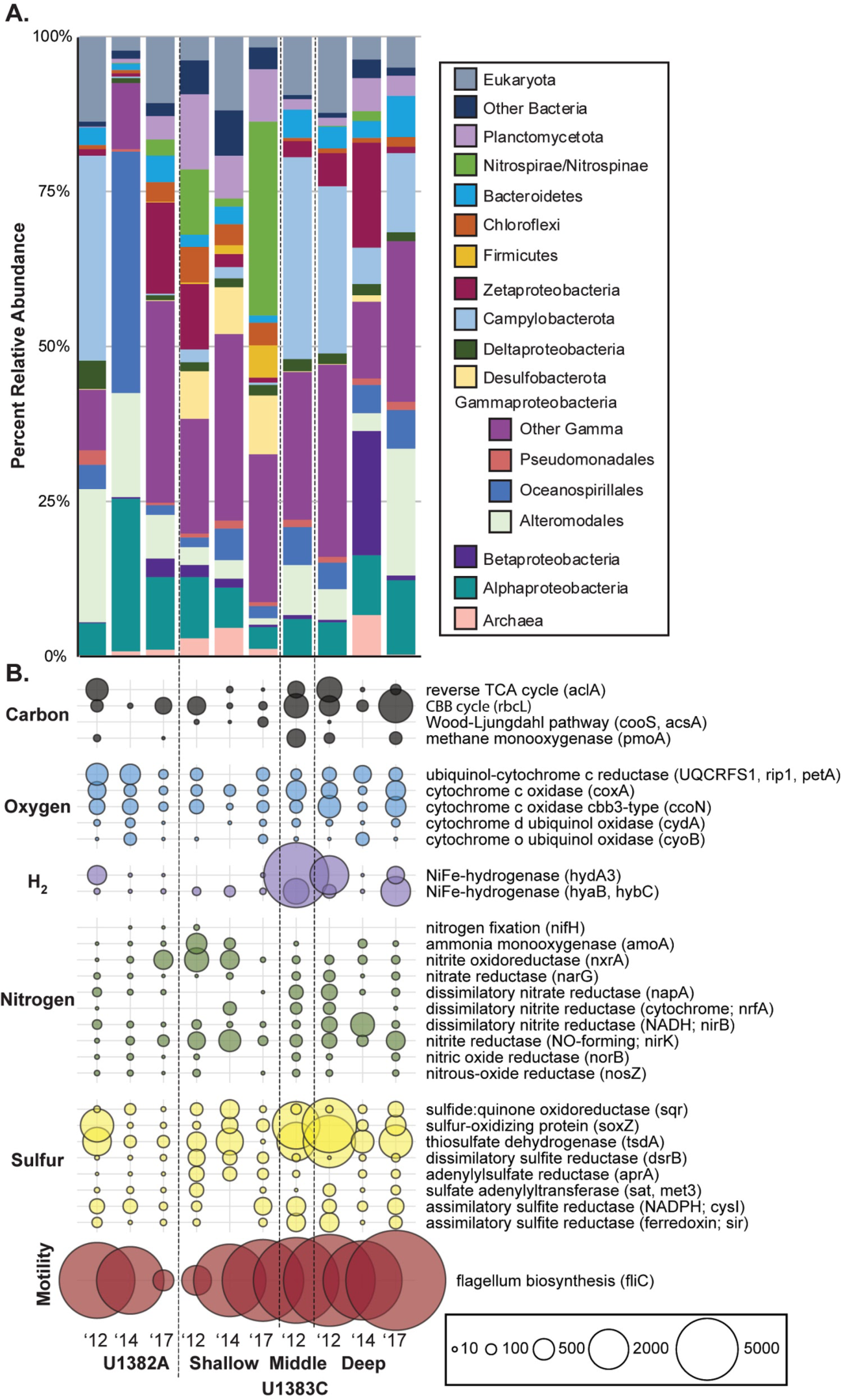
A) Relative abundance of taxa associated with the small subunit (16S/18S) and large subunit (23S/28S) ribosomal genes annotated in the metatranscriptomes. Ribosomal genes identified using SortMeRNA and annotated using UCLUST. B) Normalized abundance of transcripts of key genes for carbon, oxygen, hydrogen, nitrogen, sulfur, and flagellum biosynthesis within the metatranscriptomes at each site over three sampling years, in transcripts per million reads (TPM).

**Figure 2.**
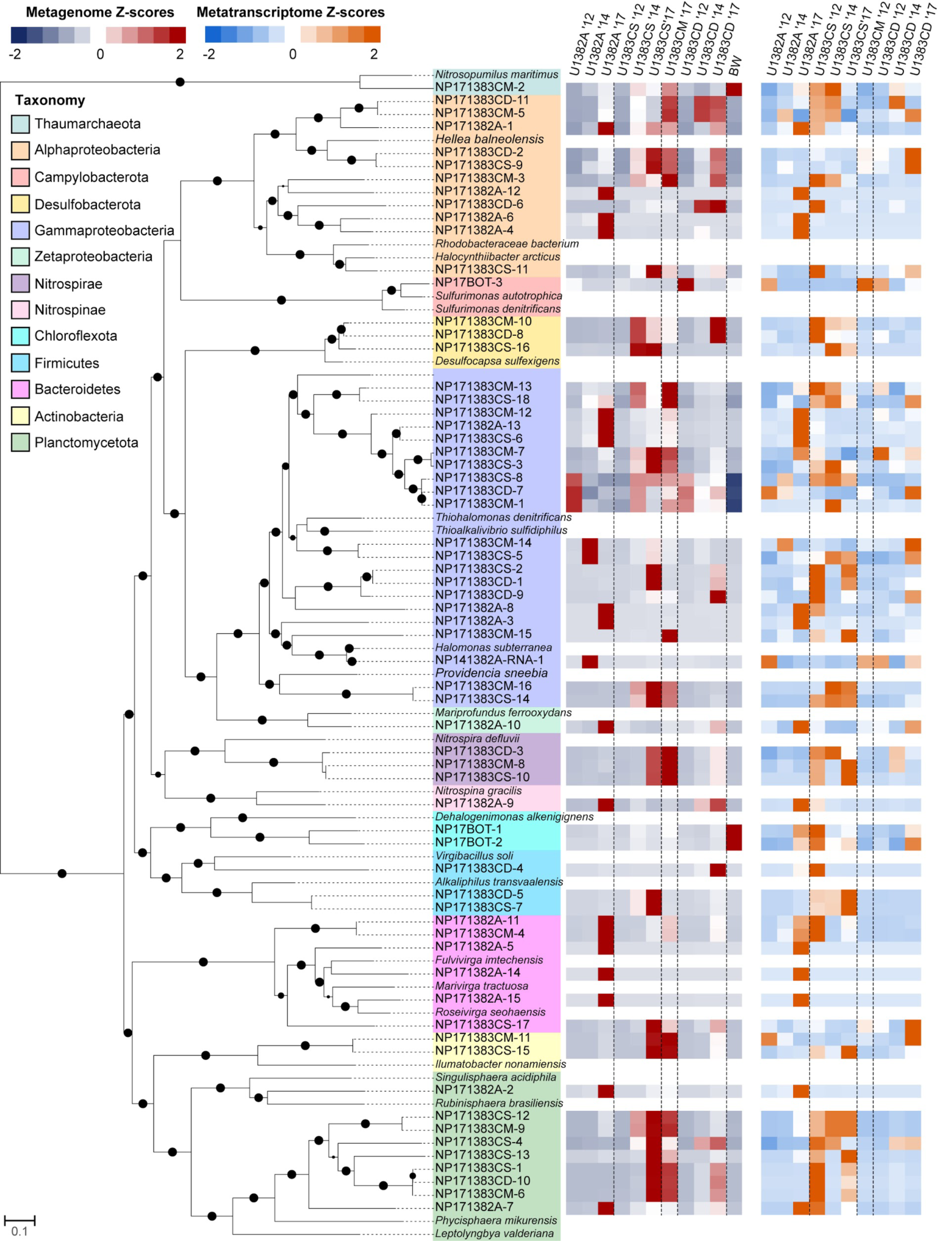
Bootstrapped phylogenetic tree of “high completion” metagenome assembled genomes (MAGs). Circles at branch notes denote bootstrapping scores (larger circles = higher scores). Includes competitive recruitment mapping of MAGs to metagenomes (red and dark blue) and metatranscriptomes (orange and blue). Relative abundance is expressed as z-scores, calculated by subtracting the average relative abundance across all samples from the MAG relative abundance, and dividing by the sample standard deviation across all samples.

**Figure 3.**
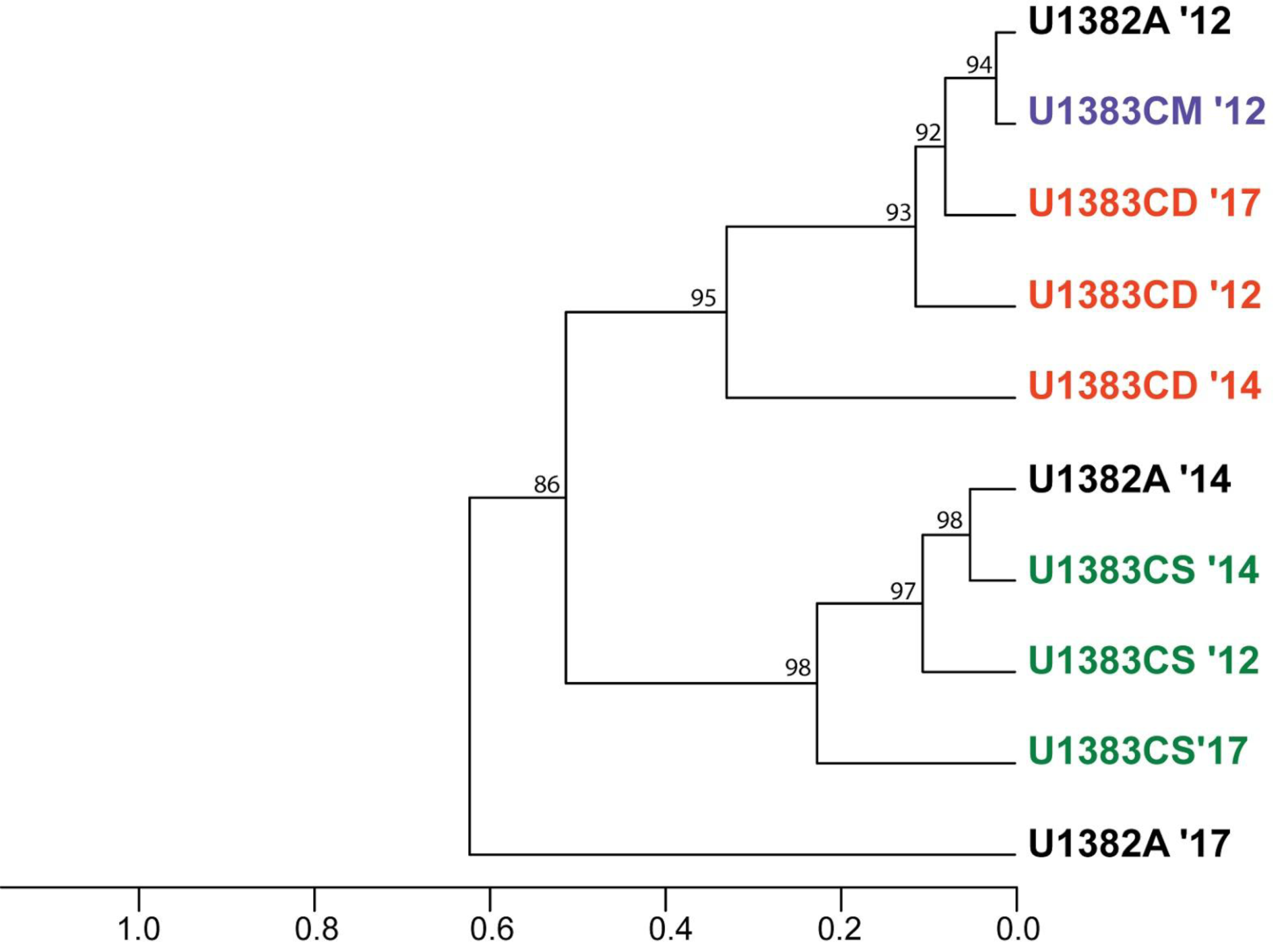
Hierarchical clustering of MAG abundances in the metatranscriptomes produced using multiscale bootstrap resampling. The scale bar indicates correlation distance between samples.

**Figure 4.**
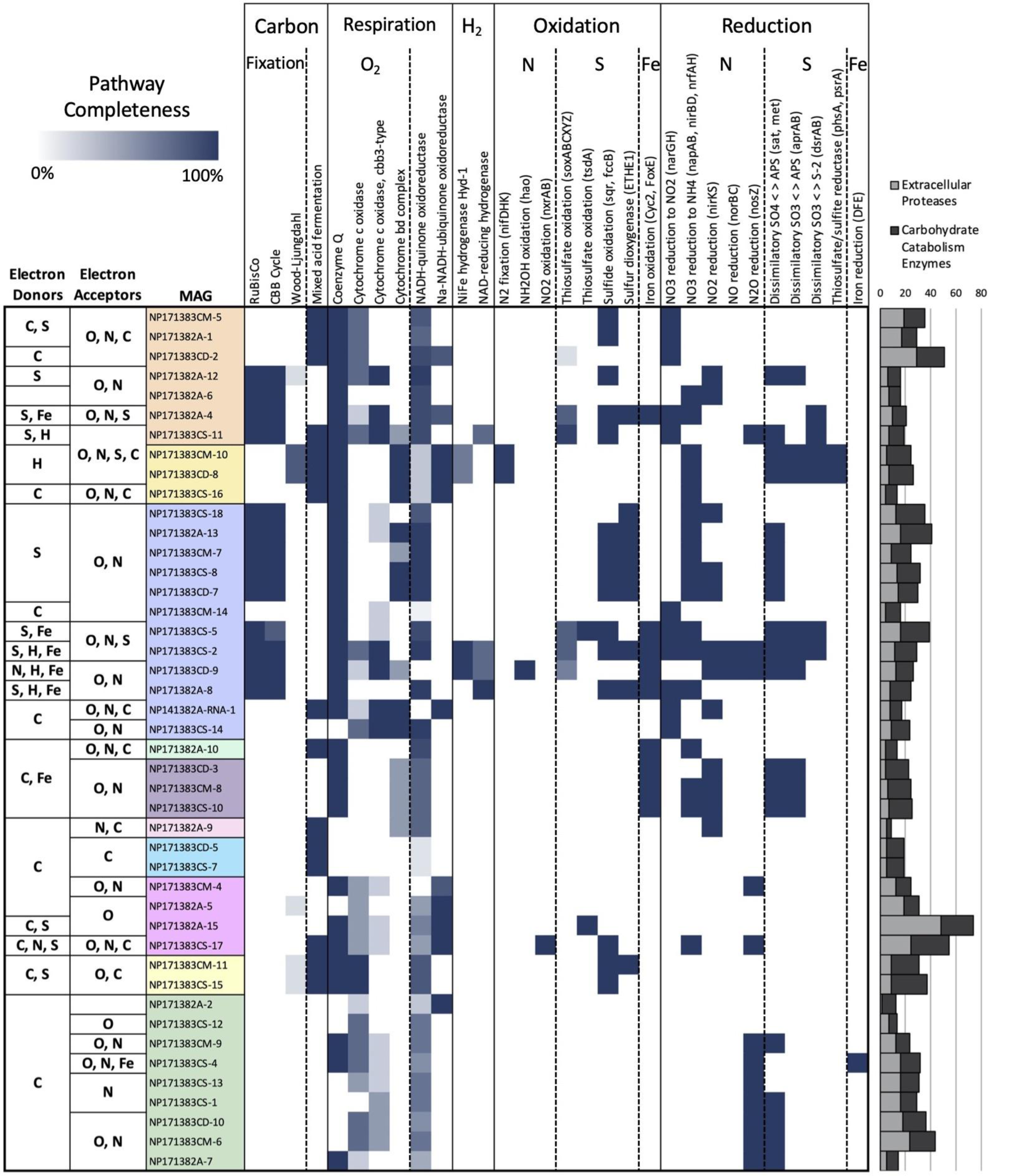
Completeness of select geochemical pathways of interest in 2017 abundant high-completion MAGs determined using KEGG-Decoder. Completeness of each enzymatic pathway is expressed as a percentage (0 to 100%). Bar graph at right depicts counts of extracellular proteases and carbohydrate-catabolism enzymes in each MAG.

**Figure 5.**
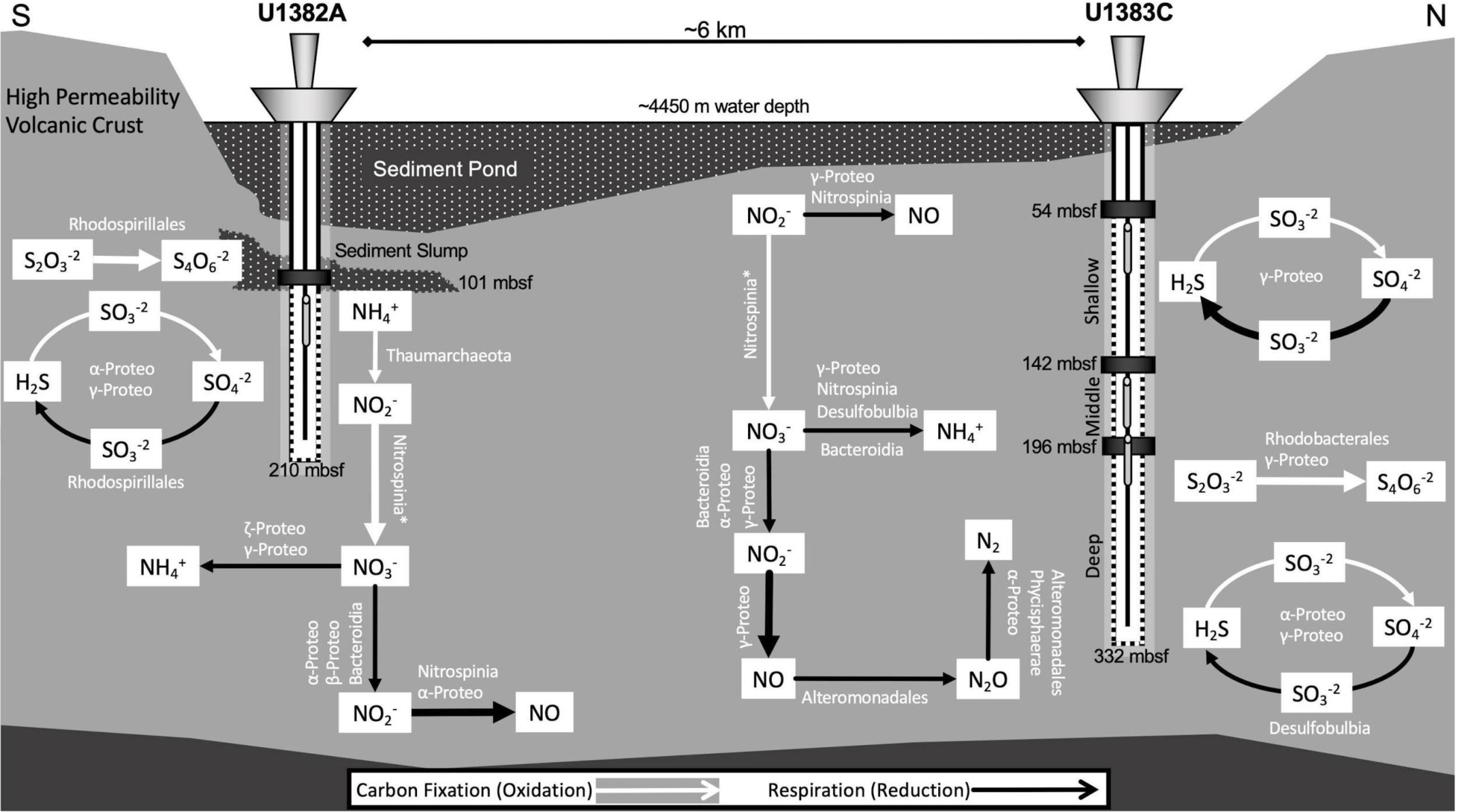
Schematic of the nutrient cycling processes occurring in North Pond, inferred from metatranscriptomic reads as mapped to MAGs, metagenome assemblies, and taxonomic databases. Pathways associated with carbon fixation are indicated with white arrows and respiration is indicated with black arrows. Thicker arrows represent processes that are particularly abundant in the metatranscriptomes. In the case where a gene was not found in a MAG, we used the taxa annotations performed by IMG (these taxa are indicated with an asterisk).

### DNA/RNA Extraction and Library Construction

DNA was extracted from 2017 North Pond samples (half of 47 mm flat) filters following a method adapted from [22, 23] (Supplemental Methods). Extracts were quantified using a Quant-iT PicoGreen dsDNA assay kit (Thermo Fisher Scientific) according to the manufacturer’s instructions, using a fluorescent plate reader. DNA concentrations of all samples are shown in Supplementary Table 1.

RNA was extracted from 47 mm flat filter halves from all three expeditions (2012, 2014, and 2017, Table 1) using a modified version of the mirVana RNA extraction kit (Ambion) protocol (Supplemental Methods). Extracts were quantified using a Quant-iT RiboGreen RNA assay kit (Thermo Fisher Scientific). One whole and one-half filter from each sampling horizon from 2017 were extracted in this manner and the extracts pooled for downstream analysis. For the 2012 and 2014 samples, one half filter was extracted. RNA concentrations are shown in Supplementary Table 2.

Metagenomic libraries were prepared from the 2017 DNA extractions using an Ovation Ultralow V2 DNA-Seq library preparation kit (NuGEN), according to the manufacturer’s instruction as described in the Supplementary Methods.

RNA yields were too low to use an RNA sequencing library preparation kit. Instead, we used a SuperScript III First-Strand Synthesis System (Invitrogen), followed by an NEBNext Ultra II Non-Directional RNA Second Strand Synthesis Module (New England Biolabs), to produce double-stranded cDNA from our RNA extracts. The cDNA was then purified using a MinElute PCR Purification Kit (QIAGEN), and libraries were prepared as for DNA, as described in the Supplementary Methods.

Metagenomes from 2017 and metatranscriptomes from 2012, 2014, and 2017 were sequenced on an Illumina NextSeq 500 at the W.M. Keck sequencing facility at the Marine Biological Laboratory, resulting in an average read length of 151 bp.

### Metagenome Assembly and Mapping

The 2017 metagenomes were sequenced twice due to poor clustering on the first run. Raw sequence data from both runs was visualized using fastqc [24] and quality filtered using Minoche et al. [25] quality filtering scripts [26]. Filtered R1 and R2 reads from each run were interleaved using fq2fa (https://pypi.org/project/fq2fa/) and the interleaved files from the two runs were concatenated for all downstream processing. Assemblies were constructed using IDBA-UD version 1.1.3 [27] and quality assessed with MetaQUAST v5.0.2 [28]. Quality filtered metagenomic reads from 2012 and 2014 were reassembled using the same pipeline. Total number of reads, quality filtering results, and assembly stats are reported in Supplementary Table 2.

Metagenome assemblies were uploaded to the Joint Genome Institute’s Integrated Microbial Genomes and Microbiomes (IMG/M) system [29] for annotation using default IMG parameters [30]. ORF information was extracted from the resulting general feature format (.gff) and fasta files using gff2seqfeatures.py (https://github.com/ctSkennerton/scriptShed). The resulting fasta feature nucleotides (.ffn) files were indexed in Kallisto [31], and the R1 and R2 quality filtered read files from each metagenome were mapped to a concatenated fasta feature nucleotides (.ffn) file of all metagenome ORFs to normalize and quantify gene abundance in transcripts per million reads (TPM), using kallisto quant (default parameters). Because the KEGG database does not differentiate between *nxrA* and *narG* or *pmoA* and *amoA* genes as they are highly orthologous, we discerned nitrite oxidation from nitrate reduction and methane oxidation from ammonia oxidation as described in supplementary methods.

rRNA was removed from the metagenomes by mapping the reads against the SILVA small subunit (SSU; 16S/18S) and large subunit (LSU; 23S/28S) ribosomal databases (release 132 [32]) using SortMeRNA (default parameters [33]). Taxonomy was assigned to the rRNA reads using assign_taxonomy.py in QIIME [34] with the default assignment method UCLUST [35] against the SILVA SSU and LSU databases.

### Metatranscriptome Assembly and Mapping

Quality filtering and assembly were performed on the metatranscriptomes in the same manner as for the 2017 metagenomes (Supplementary Methods). rRNA was removed from the quality-filtered metatranscriptome reads using SortMeRNA and taxonomy was assigned to the rRNA files using the SILVA SSU/LSU databases and QIIME as above. The metatranscriptomes with rRNA removed were mapped to a concatenated fasta feature nucleotides (.ffn) file of all metagenome ORFs in Kallisto.

### Binning and Metagenome Assembled Genomes (MAGs)

Assembled metagenomes from 2017 and metatranscriptomes from 2012, 2014, and 2017 were binned with Binsanity [36] and bin quality was assessed using CheckM version 1.0.11 [37] (Supplemental Methods). High-completion bins (Supplementary Table 3), hereafter referred to as metagenome-assembled genomes (MAGs), were taxonomically classified using Phylosift [38] and GTDB-Tk [39]. The MAGs were then imported into KBase [40] and a phylogenetic tree with 100 bootstrap replicates was made using the SpeciesTreeBuilder application with FastTree2 [41]. Tree formatting was performed using the Interactive Tree of Life [42]. Functional orthologies as defined by the KEGG database [43] were annotated using MetaSanity [44] (Supplemental Methods). Genes related to iron acquisition, storage, and oxidation/reduction were annotated using the FeGenie tool [45].

### Mapping MAGs to Metagenomes and Metatranscriptomes

The relative abundance of each MAG in the metagenomes and metatranscriptomes was calculated using a competitive recruitment approach (Supplemental Methods). Z-scores of the normalized relative fractions were calculated using the average and sample standard deviation of the relative fraction of each MAG across all metagenomes and across all metatranscriptomes.

The metagenome and metatranscriptome MAG mapping data was also used to generate dendrograms using the pvclust package in R [46]. Relative fraction data was scaled using z-scoring, p-values were calculated using an unweighted pair group method with arithmetic mean for hierarchical clustering, and distances between samples determined using correlation as a measure of similarity. Dendrograms were then generated with 1000 bootstrap replications.

### Data Availability

Data from 2017 metagenomes and 2012, 2014, and 2017 metatranscriptomes has been deposited at SRA under the BioProject accession numbers PRJNA554681 and PRJNA522799, respectively, and raw sequence reads are available with accession numbers SRR9705319-SRR9705323, SRR8590998-SRR8591005, and SRR10499325-SRR10499326. Assembled contigs for each 2017 metagenomic library are publicly available via IMG/MER under submission numbers 3300029216 and 3300029223-3300029226. Contigs for each individual 2017 MAG are available through FigShare at https://figshare.com/articles/dataset/North_Pond_2017_High_Completion_Bins/12389789. Raw sequence reads from Meyer et al. [9] constituting the metagenomic samples from 2012, are available under the BioProject accession no. PRJNA280201. Data from Tully et al. [13], including the 2013 and 2014 metagenomes, are available under the BioProject accession no. PRJNA391950 and raw sequence reads are available with accession no. SRX3143886-SRX3143902. Reassembled 2012-2014 metagenomes used in this study are available via IMG/MER under submission numbers 3300029924-3300029927 and 3300029949-3300029950. All accession numbers are shown in Supplementary Table 2.

## Results

While DNA and RNA yields were generally low (<1 ng/µl, Supplementary Table 1), consistent with cell quantification (Table 1), we were successful in generating the first metatranscriptomes from this environment. Together, we examined over one billion high-quality paired-end Illumina sequencing reads from 11 metagenomes and 10 metatranscriptomes over three sampling periods spanning five years from four different subseafloor fluid horizons and deep bottom water at the North Pond site (Table 1, Supplementary Table 3).

While similar taxa were present across all metagenomic and metatranscriptomic samples, their relative abundances fluctuated over time. Most samples were dominated by Gamma- and Alphaproteobacteria, as well as Bacteroidetes and Zetaproteobacteria (Figure 1A; Supplementary Figure 1A). Campylobacterota represented a large proportion of the total microbial community in 2012, but in 2017 they made up a much smaller percentage of the crustal fluid community (Figure 1A), though they remained a significant taxonomic group in the 2017 bottom water (Supplementary Figure 1A). Additionally, Nitrospirae and Nitrospinae assigned reads, which represented ≤3% of each metagenome (Supplementary Figure 1A), composed 31% of the metatranscriptome-mapped rRNA reads in U1383C Shallow in 2017 (Figure 1A). Firmicutes-assigned reads (chiefly family *Clostridiaceae*) accounted for >50% of the mapped rRNA reads in the 2017 U1383C Shallow metagenome, but represented only 5% of the mapped metatranscriptome rRNA reads in U1383C Shallow in 2017 (Figure 1A).

### Metabolic, motility, biofilm, and stress transcript abundance across samples

To examine functional potential at the community level across samples, we examined genes encoding the active sites of key enzymes for cycling carbon, hydrogen, oxygen, nitrogen, and sulfur. Overall, these genes were equally abundant in the metagenomes across sampling times and depth horizons (Supplementary Figure 1B), while transcript abundance was much more variable (Figure 1B).

In most samples, carbon fixation via the Calvin-Benson-Bassham (CBB) cycle was more abundant than the reverse tricarboxylic acid cycle (rTCA), as indicated by the number of transcripts for the large subunit of RuBisCo (*rcbL*) (∼10-1400 tpm) versus the ATP citrate lyase alpha-subunit (*aclA*) (∼0-300 tpm), (Figure 1B). In 2017, *rcbL* and *aclA* transcripts were highest in U1382A and U1383C Deep than in U1383C Shallow by 1-2 orders of magnitude (Figure 1B). The carbon-monoxide dehydrogenase catalytic subunit (*cooS*) of the Wood-Ljungdahl pathway was transcribed in U1383C Shallow in all years and U1383C Deep in 2012 (∼1-80 tpm). Aerobic oxidation of methane (*pmoA*) transcripts were observed in 2012 and 2017 but not 2014 (Figure 1B).

Genes for aerobic oxidative phosphorylation (cytochrome c oxidase (*coxA*), cytochrome o ubiquinol oxidase (*cyoB*), and ubiquinol-cytochrome c reductase (*UQCRFS1, rip1, petA*)) and cytochromes related to microaerophily and low oxygen tolerance (*cbb*_*3*_-type cytochrome c oxidase (*ccoN*) and cytochrome d ubiquinol oxidase (*cydA*)) were transcribed in every sample (Figure 1B). *cbb*_*3*_-type cytochrome c oxidases were 3-7x more abundant in 1383C Deep than in U1382A or 1383C Shallow. NiFe-hydrogenase transcripts used for anaerobic respiration and oxidative stress response (*hyaB, hybC, hydA3*) were highest in U1383C Middle in 2012 and U1383C Deep in 2012 and 2017, and were 30-1000x more abundant in U1383C Deep than in U1382A or 1383C Shallow in 2017 (Figure 1B).

Among transcripts involved with nitrogen cycling, the most abundant were associated with ammonia oxidation, nitrate reduction, and nitrite reduction. Transcripts for *amoA* were most prevalent in U1383C Shallow in 2012 (∼465 tpm) and 2014 (∼110 tpm), but in 2017 were only found in U1382A and U1383C Deep, at an order of magnitude lower abundance (∼10-15 tpm) (Figure 1B). In the metagenomes, *amoA* genes displayed the highest counts in the 2017 bottom seawater sample (Supplementary Figure 1B). The catalytic subunit of nitrite oxidoreductase (*nxrA*) was transcribed in every sample; transcripts were highest in U1383C Shallow in 2012 (∼660 tpm) and 2014 (∼380 tpm) and U1382A in 2017 (∼390 tpm) (Figure 1B). Nitrate reduction through either denitrification (*narG*) or the dissimilatory reduction of nitrate to ammonia (DNRA; *napA*) was also transcribed in every sample ranging from 0.5 to >250 tpm, but these processes were most abundant in 2012, in U1383C Middle and Deep (Figure 1B). Nitrite reduction via both DNRA (*nirB*) and denitrification (*nirK*) was more transcribed than nitrate reduction across samples and years, ranging from ∼10-640 tpm (Figure 1B). Nitric oxide reduction to N_2_ was transcribed primarily in samples taken in 2012, but was also observed in U1383C Deep in 2017 (Figure 1B). Very few transcripts for nitrogen fixation (*nifH*) were detected. Nitrogen reduction genes were more abundant than nitrogen oxidation genes in the metagenomes across all samples and years (Supplementary Figure 1B).

Genes for the oxidation or assimilation of sulfur were more transcribed (by 1-2 orders of magnitude) than for dissimilatory sulfate reduction. The most transcribed sulfur cycling genes were those involved in thiosulfate oxidation: thiosulfate oxidase (*soxZ*, up to ∼3800 tpm) and thiosulfate dehydrogenase (*tsdA*, up to ∼3500 tpm) (Figure 1B). While dissimilatory sulfate reduction gene transcripts (*dsrB, aprA, sat, met3*) were present in every sample, they were highest in U1383C Shallow across all sampling years (∼100-400 tpm) (Figure 1B).

Transcripts associated with chemotaxis and flagellum biosynthesis were far more abundant in the metatranscriptomes than those associated with biofilm formation (Supplementary Figure 3). Flagellin (*fliC*) was among the most abundant transcripts in every sample, and was in the top three most abundant transcripts in every 2017 sample (Figure 1B, Supplementary Table 4). The top three most abundant transcripts in U1382A and U1383C Deep in 2017 also included *pilA* (type IV pilus assembly protein PilA), associated with twitching motility (Supplementary Figure 3, Supplementary Table 4). Transcripts for catalases associated with oxidative stress (*katG, katE, CAT, catB, srpA, ahpC*), cold shock, and phage shock proteins were also high in 2017 in U1383C Deep, but displayed far lower transcription levels in other samples (Supplementary Figure 3; Supplementary Table 4).

### Metagenome-Assembled Genomes (MAGs)

A total of 449 MAGs were obtained from the 2017 metagenomes and 2012, 2014, and 2017 metatranscriptomes using BinSanity. Two bins were acquired from the metatranscriptomes, both from the 2014 U1382A sample. 64 non-redundant, high-quality, high-completion bins (hereafter referred to as metagenome-assembled genomes, or MAGs) were chosen for downstream analyses (Supplementary Table 3). MAGs were designated “NP” (for “North Pond”), followed by the 2-digit sampling year (“17”), followed by the sampling location (e.g. 1382A).

The taxonomic identities of these MAGs closely matched the taxonomic annotations of the 16S/23S rRNA gene mapping in the metagenomes and metatranscriptomes (Figure 1, Supplementary Table 3). More than half (33) of the MAGs belonged to the Proteobacteria: Gammaproteobacteria (21), Alphaproteobacteria (11), and Zetaproteobacteria (1). Other MAGs were annotated as Planctomycetota (9), Bacteroidetes (6), Desulfobacterota (3), Firmicutes (3), Nitrospirae (3), Actinobacteriota (2), Chloroflexota (2), Campylobacterota (1), and Nitrospinae (1). One archaeal MAG was also identified and classified as a member of the Thaumarchaeota.

We compared the relative abundance of each high-completion MAG across all metagenome and metatranscriptome samples (Figure 2). The most abundant MAGs in the metagenomes were generally observed in North Pond samples from 2017, while some MAGs were found in multiple fluid horizons and years. Transcriptome reads mapping to MAGs correlated with metagenome reads mapping to MAGs in some cases, and hierarchical clustering of MAG transcript abundances showed that the U1383C Middle and Deep horizon samples clustered together, and the U1383C Shallow samples clustered together (Figure 3). Clustering of MAG abundances from the metagenomes indicated all 2012 samples were most similar to one another (Supplementary Figure 4). U1382A 2017 was distinct from other samples in both the MAG metatranscriptomes (Figure 3) and metagenomes (Supplementary Figure 4).

Twenty-two MAGs contained complete or mostly complete (>80%) carbon fixation pathways (Figure 4, Supplementary Table 5). Most of the MAGs with the potential for carbon fixation contained pathways for the oxidation of sulfide, thiosulfate, or sulfite, making sulfur compounds a probable electron donor (Figure 4, Supplemental Figure 5). Three MAGs contained genes associated with the oxidation of nitrogen compounds, but only one had the capability to fix carbon (Figure 4, Supplementary Table 5). Nine MAGs possessed genes for the oxidation of ferrous iron and five of these, one Alphaproteobacteria and four Gammaproteobacteria, also contained the genes for RuBisCo and the CBB cycle (Figure 4). One Alphaproteobacteria and two Gammaproteobacteria contained genes for NiFe and NAD-reducing hydrogenases (Figure 4). The majority of MAGs with carbon fixation pathways also contained numerous extracellular protease and carbohydrate catabolism genes (Figure 4, Supplementary Tables 6 and 7). The remaining 42 MAGs lacked carbon fixation genes, but several had the ability to oxidize sulfur compounds and/or ferrous iron (Figure 4, Supplementary Table 5).

While the majority (52) of the MAGs contained cytochromes for aerobic respiration (Supplementary Table 5), all but six aerobic MAGs also had the capability to use other electron acceptors (nitrogen or sulfur) or contained fermentative pathways (Figure 4, Supplementary Table 5). All of the MAGs that contained carbon fixation and sulfur oxidation pathways had the ability to use either oxygen or oxidized nitrogen compounds as electron acceptors (Figure 4, Supplementary Table 5).

## Discussion

Low-temperature, off-axis environments represent the majority of global hydrothermal fluid circulation in the ocean, but the microbial communities in the subseafloor of young ridge flank oceanic crust are sparsely sampled or understood in comparison to other subseafloor habitats such as marine sediments. The goal of this study was to reconstruct microbial metabolic potential, transcript abundance, and community dynamics in the crustal fluids of the cool, oxic ridge flank North Pond using metagenomics and metatranscriptomics. This work represents the first metatranscriptomic data recovered from the cool, oxic subseafloor aquifer, where low biomass presents significant challenges to microbial studies.

Accessing the crustal aquifer in sedimented regions requires ocean drilling and observatory CORK installations to enable collection of crustal fluids, thus all results must be interpreted in the context of the disturbance caused by drilling and subsequent recovery from these operations. Time series sampling at North Pond from 2012-2017 has therefore been critical to resolving when the crustal aquifer returned to its pre-drilling state. Geochemical measurements taken over this 6-year period indicate the fluid chemistry rebounded by 2017 [12], and cell count data in 2017 are the lowest since the CORKs were first sampled (Table 1, Trembath-Reichert et al. In Review). Geochemical and heat-flow data also show that U1382A experiences greater connectivity to the open ocean than U1383C, although all fluids are geochemically very similar to deep seawater [9, 47]. Heat-flow data [48, 49] and tracer experiments [12] indicate rapid lateral fluid flow from U1382A to U1383C through the porous crust, while radiocarbon data suggests potential residence times on the order of hundreds (U1382A) or thousands (U1383C) of years [10]. Within these important geochemical and hydrogeochemical contexts, we therefore focused our analyses on understanding the microbial community in 2017, the samples furthest removed from the disturbances caused by drilling, while also allowing us to examine how the community has changed over time by comparing 2017 samples to those collected in 2012-2014 [9, 13].

Analysis of metagenomic data from North Pond samples in 2012-2014 suggested a high degree of functional redundancy despite differences in community membership across samples and years [13]. The addition of 2017 metagenomic data as well as time series metatranscriptomic data broadly supports this finding and shows that while most examined functional genes are present across many samples and taxa (Supplementary Figure 1B), their transcript abundance varies with both time and location within the crust (Figure 1B). Our analysis via the presence and transcription of metagenome-assembled genomes further indicates that while some members of the community are present and transcribed in multiple years and sampling horizons, many were only abundant and active in 2017 samples (Figure 2). For example, many MAGs at U1383C Shallow were transcribed across all years in that horizon, but most MAGs at U1382A and U1383C Deep were only transcribed in 2017, with few shared members between these two locations, consistent with heat-flow and geochemical data. Hierarchical clustering of MAG abundances in the metatranscriptomes suggests that overall, the same MAGs were active in U1383C Shallow and in U1383C Deep across all sampling years, while U1382A was more variable (Figure 3), which may reflect greater influence of sediment interactions at this sampling horizon.

Thus, based on mapping 2017 transcripts to annotated metagenome ORFs, the taxa identified in the metatranscriptomes, and the predicted metabolisms of the most abundantly transcribed MAGs, as well as building on previous work at North Pond, we constructed a conceptual model for the key carbon, nitrogen, and sulfur cycling reactions most likely occurring in the North Pond crustal fluids in 2017 and the microbial taxa which carry out these reactions (Figure 5). We distinguished reactions that are connected to carbon fixation (oxidation of nitrogen or sulfur) and respiration (reduction of nitrogen or sulfur). The model is a simplified representation of the most abundant microbially-mediated processes occurring at different locations and depths in the North Pond crustal aquifer.

Our analyses also indicated a number of lifestyle strategies and adaptation of microbial communities in the crustal fluids. For example, the abundance of transcripts associated with chemotaxis and flagellar motility, and the concurrent lack of transcripts for biofilm formation, suggest that the microbial community captured in these samples is motile, as biofilm formation is generally preceded by inhibition of flagellar gene transcription [51, 52, 53, 54]. A prevalence of motility and chemotaxis genes has been described in warm, anoxic crustal fluids in Juan de Fuca Ridge flank [55]. Additionally, we found evidence for several stress response mechanisms which were highly transcribed in U1383C Deep in 2017, including oxidative stress response, survival under nutrient- and energy-limited conditions (phage shock protein A) [56], and cold shock (CspA) [57] (Supplementary Figure 3, Supplementary Table 4). These results suggest that especially at greater depth in the basement at North Pond, microbial communities experience multiple stressors that may be linked to energy and temperature fluctuations.

### Carbon Fixation and Turnover

The microbial community of the North Pond aquifer appears to be predominantly heterotrophic or mixotrophic (Figure 4, Supplementary Table 5), with carbon fixation primarily occurring in U1382A and U1383C Deep (Figure 1B). MAGs that possessed both carbon fixation and organic carbon degradation genes were highly expressed in both the U1383C Shallow and U1383C Deep metatranscriptomes (Figure 2, Figure 4). These mixotrophs may perform heterotrophy in U1383C Shallow, and carbon fixation in the Deep horizon. These observations are consistent with radiocarbon data from North Pond, which suggests rapid turnover of semi-labile DOC sourced from the deep ocean, followed by slower removal of more refractory components of the DOC pool [10]. Chemosynthetic DOC produced in situ is likely turned over rapidly in the shallow crustal fluids, but Δ^14^C values in the Deep horizon of U1383C suggest a net contribution of ^14^C-enriched DOC from chemoautotrophy [10]. Data from 2017 bulk and single-cell rate measurements (Trembath-Reichert et al. In Review) likewise found a preference for uptake of organic substrates over DIC under in situ conditions. However, these experiments also showed evidence for biomass production from DIC, and rates of activity varied over 8 orders of magnitude for both inorganic and organic carbon uptake, consistent with the mixotrophic genetic signatures observed in our study. Two recent microbial studies of subseafloor rocks recovered via ocean drilling also showed a dominance of heterotrophic bacteria and little evidence for autotrophic processes [58, 59], suggesting mixotrophy and heterotrophy may be a common strategy in the cool, crustal subseafloor habitat.

Fixation of carbon is most likely linked to the oxidation of sulfide and thiosulfate, as evidenced by both the metatranscriptomic (Figure 1B) and MAG data (Figure 4 [13]). Oxidation of sulfur compounds is a well-characterized source of electrons for chemolithoautotrophy in the deep sea [60, 61]. Most of the sulfur-oxidizing chemoautotrophic MAGs were capable of using more than one type of sulfur compound, including oxidation of thiosulfate (*soxZ, tsdA*), as well as sulfide:quinone oxidoreductase (*sqr*) in U1383C Deep (Figure 1B). While hydrogen sulfide has not been detected in the borehole fluids [9, 12], the oxidation of iron in sulfide complexes in the crustal rocks may allow access to both sulfide [62] and thiosulfate [63] for carbon fixation.

Previous metagenomic work at North Pond suggested the potential for some microbes to use H_2_ and Fe+2 to drive biomass production [13], by using the redox gradient between reduced material in basalt and oxygenated aquifer fluids [64, 65]. Oxidation of ferrous iron has been described in a low temperature deep-sea environment near the Juan de Fuca hydrothermal field [66]. Our results indicate that some members of the North Pond crustal fluid microbial community may use hydrogen and/or ferrous iron as electron donors for carbon fixation. We identified six MAGs that contained the CBB cycle and a NiFe hydrogenase and/or the iron oxidation gene Cyc2 (Figure 4), an outer-membrane cytochrome that has been characterized in multiple iron-oxidizing bacterial lineages [67]. NiFe-hydrogenases catalyze the conversion of H_2_ to protons and electrons and are generally inhibited by oxygen, but oxygen-tolerant and aerobic NiFe hydrogenases have been described in marine Gammaproteobacteria [68, 69]. We also identified three MAGs with high transcription in U1383C Shallow and high abundance in the U1383C Deep 2017 metagenome that were taxonomically assigned as *Desulfocapsa* (*Desulfobulbaceae*), two of which contained NiFe hydrogenases (Figure 4; Supplementary Table 3). Members of this clade simultaneously oxidize and reduce sulfur compounds for energy while using electrons from H_2_ to fix carbon via the Wood-Ljungdahl pathway [70]. Transcripts of the carbon-monoxide dehydrogenase catalytic subunit (*cooS/acsA*) were present in the U1383C Shallow metatranscriptome in 2017 (Figure 1B), on contigs annotated as *Desulfobulbaceae* (Supplementary Table 4).

### Respiration, Oxygen, and Anaerobiosis

MAGs possessing carbon fixation pathways all contained at least one aerobic or microaerophilic terminal oxidase (Figure 4), suggesting that carbon fixers in the aquifer use oxygen as a terminal electron acceptor. However, if oxygen is limiting or unavailable, nitrate or nitrite may be used as an alternative (Figure 4, [13]). The coupling of sulfur, sulfide, and thiosulfate oxidation to denitrification is well documented in multiple bacterial clades (e.g., [71, 72]). Anaerobic oxidation of sulfur [73] and sulfide [74] coupled to DNRA has also been described in marine environments.

The complete suite of transcripts for denitrification from NO_3_ to N_2_ was identified in U1382A and U1383C Deep (Figure 1B). However, transcripts for the final two steps of denitrification were not abundant in U1382A (<2 tpm; Figure 1B), and none of the abundant MAGs in U1382A were capable of reduction of NO to N_2_O (Figure 2, Figure 4). Reduction of nitrogen species from NO to N_2_ therefore appears to occur primarily in U1383C Deep, carried out by the Alphaproteobacteria, Gammaproteobacteria (Alteromonadales), and Planctomycetota (Phycisphaerales) (Figure 5). The majority of MAGs that contained denitrification genes also contained aerobic terminal oxidases, except for two of the Planctomycetota (Figure 4). While denitrification is generally a strictly anaerobic process, aerobic denitrifying bacteria have been described [75, 76].

Sulfur reduction was a less common anaerobic respiration strategy, both in the metatranscriptomic data (Figure 1B) and in the MAGs (Figure 4). Six MAGs in total had the capability to reduce sulfite to sulfide, all of which contained pathways for carbon fixation: 2 Desulfobacterota, 2 Alphaproteobacteria, and 2 Gammaproteobacteria. (Figure 4). One MAG, a Gammproteobacteria (NP171383CS-2), contained a complete pathway for the reduction of dimethyl sulfoxide (Supplementary Table 5).

Anaerobic processes appear to be most relevant in U1383C Deep, based on the presence of denitrification transcripts and denitrifying MAGs, *cbb*_*3*_-type cytochrome c oxidases, NiFe hydrogenases, and catalases associated with oxidative stress. This is consistent with the lower (173 µM [12]) oxygen concentrations present in this horizon compared to other horizons [12, 13]. Additionally, samples collected in 2012, 6 months after borehole drilling ceased, included particle-laden fluids which have not been observed since [9, 12]. Organic aggregates provide microenvironments of anoxia in otherwise oxic seawater, vastly expanding the available niche space of denitrifying and sulfate-reducing bacteria in the global ocean [50]. This may explain why we observed high abundances of NiFe-hydrogenase transcripts associated with anaerobic metabolisms in 2012, and why abundances of these genes were mostly confined to the Deep horizon of U1383C in 2017, post-drilling recovery (Figure 1B). It is possible that the crustal environment may contain slower flow paths or stagnant zones where oxygen becomes depleted, allowing for the anaerobic metabolisms we observed to take place. Furthermore, the abundance of motility and flagellar genes in the metatranscriptomes and the lack of genes for biofilm formation (Supplementary Figure 3) also suggests that the microbial community captured in these samples is highly mobile and capable of seeking out organic particles to colonize and consume. Fractures in the basaltic basement may host more sedentary, surface-associated microbial communities with metabolisms not represented in the data presented here.

### Nitrification

Cross-hole tracer experiments during the most recent cruise to North Pond detected an increase in nitrate concentrations relative to bottom seawater, most likely the result of microbial nitrification [12]: the oxidation of ammonia to nitrite and nitrite to nitrate. Ammonium concentrations in borehole fluids are usually low (<0.1 µM ammonium), and metatranscriptomic evidence for nitrogen fixation to ammonia was lacking in the aquifer (Figure 1B). Dissimilatory nitrate reduction to ammonia (DNRA) transcripts, however, were present in all three 2017 samples (Figure 1B, Figure 5) and the DNRA pathway was present in 23 of the 64 MAGs (Figure 4). Ammonia produced by this pathway may be rapidly consumed by nitrifying archaea and bacteria.

*amoA* transcripts were present in all of the 2017 samples (Figure 1B), and were found exclusively on Thaumarchaeota-annotated contigs. We did not detect the 4-hydroxybutyrate/3-hydroxypropionate cycle used by Thaumarchaeota for carbon fixation [77] in any of our samples. However, culturing, stable isotope, and MAG data suggest many Thaumarchaeota may be mixotrophic or heterotrophic [78, 79, 80, 81, 82].

Marine nitrite oxidation is generally associated with Nitrospira and Nitrospinia bacteria, which may grow autotrophically or mixotrophically [83, 84, 85, 86]. While these phyla were most abundant in the 2017 U1383C Shallow metatranscriptome (Figure 1A), *nxrA* transcript abundance was relatively low in this sample (11-15 tpm; Figure 1B). *nxrA* transcripts were most abundant in U1382A in 2017 (Figure 1B), and associated with unbinned contigs that could broadly be assigned to the Nitrospinia (Supplementary Table 4). While we obtained one Nitrospinia and three Nitrospira MAGs, all of them were missing *nxrA* (Figure 4, Supplementary Table 3). Based on 16S/23S data and the taxonomic annotation of the metatranscriptomic contigs, we assume that oxidation of nitrite to nitrate is being carried out primarily by Nitrospinia in U1382A and U1383C Shallow (Figure 5).

Together, our results show that the microbial community in the North Pond crustal aquifer is populated by motile mixotrophic and heterotrophic bacteria that are active under both oxic and anoxic conditions. This snapshot of subseafloor microbes at North Pond represents the fluids in their most pristine state, furthest removed from the disturbances caused by drilling. While low biomass presents significant challenges for such studies, this first examination of transcripts from the cool, oxic subseafloor aquifer highlights the spatial heterogeneity of life in such fluids and the ability of microbes to respond and adapt to different regimes in the crustal matrix.

## Acknowledgements

We thank the captain and crew of the R/V Atlantis, the pilots and engineers of the ROV Jason II, and B. Orcutt, C. G. Wheat, P. Girguis, S. Shah Walter, M. Mullis, K. Yoshimora, O. Nigro, G. Stewart, K. Freel, Tim D’Angelo, C. Sullivan, and M. Rappé for their work in accomplishing the field program. The Gordon and Betty Moore Foundation sponsored most of the observatory components at North Pond through grant GBMF1609. This work was supported by NSF OCE-1745589 and OCE-1635208 to J.A.H. E.T.R. was supported by a NASA Postdoctoral Fellowship with the NASA Astrobiology Institute and a L’Oréal USA For Women in Science Fellowship. The Center for Dark Energy Biosphere Investigations (C-DEBI OCE-0939564) also supported the participation of J.A.H. and B.T. This is C-DEBI contribution number XXX.

## Notes

https://figshare.com/articles/dataset/North_Pond_Metatranscriptomics_Supplemental_Tables/12756248

https://figshare.com/articles/dataset/North_Pond_2017_High_Completion_Bins/12389789

### Competing Interest Statement

The authors have declared no competing interest.

