## Supplementary Methods and Figures for "Time-series transcriptomics from cold, oxic subseafloor crustal fluids reveals a motile, mixotrophic microbial community"

*DNA Extraction and Library Preparation*

RNAlater™ solution was removed via centrifugation and filters were washed with phosphate-buffered solution. Cells were lysed in DNA extraction buffer with a combination of freeze-thawing, Proteinase-K, lysozyme, and sodium dodecyl sulfate. DNA was purified with phenol:chloroform:isoamyl alcohol (25:24:1, pH 8.0) and precipitated with 100% isopropanol at room temperature. The DNA pellet was washed twice with ice cold 70% ethanol, dried, and resuspended in 50 µl nuclease-free water. (See [22, 23] for complete extraction protocol.)

DNA was sheared to 400 bp using a Covaris focused-ultrasonicator, and purified using Agencourt AMPure magnetic beads (Beckman Coulter) using the Ovation kit protocol. End repair, ligation, and amplification (14 cycles) were carried out according to the kit manufacturer's instructions, using nuclease-free water to elute DNA from the Agencourt beads. Prepared libraries were quantified using a DNA1000 Bioanalyzer chip (Agilent). A KAPA Library Amplification Kit (Illumina) was used to further amplify libraries with low yields (1-3 cycles). DNA fragments between 470 and 570 bp were selected from the libraries using a Pippin Prep (Sage Science).

*RNA Extraction and Library Preparation*

Cells were lysed in mirVana RNA extraction kit lysis buffer by vortexing filters with RNA Powersoil beads for 10 minutes, followed by incubation with homogenate additive for 10 minutes at -20°C. The lysate was removed by centrifugation, and nucleic acids were purified with acid:phenol chloroform and washed with 100% ethanol. RNA was captured on a provided filter cartridge and eluted using two 50-ul aliquots of elution solution. Remaining DNA was removed using a Turbo-DNAase kit (Ambion) and extracts were cleaned with a RNeasy MinElute Cleanup kit (QIAGEN). RNA was then reverse transcribed to cDNA as described in the Methods.

cDNA was sheared to 400 bp using a Covaris focused-ultrasonicator, and purified using Agencourt AMPure magnetic beads (Beckman Coulter) using the Ovation kit protocol. End repair, ligation, and amplification (15 cycles) were carried out according to the kit manufacturer's instructions, using nuclease-free water to elute cDNA from the Agencourt beads. A KAPA Library Amplification Kit (Illumina) was used to further amplify (3 cycles) libraries with low yields. cDNA fragments between 400 and 570 bp were selected from the libraries using a Pippin Prep (Sage Science). For the U1383C Shallow (2012) and U1383C Deep (2017) samples, processing resulted in an overall smaller size distribution, and thus 200-370 bp fragments were selected for sequencing.

*Metagenome Assembly*

Assemblies were constructed using IDBA-UD version 1.1.3 [27] with a minimum contig length of 450 bp. For comparative purposes, multiple assemblies were produced, using iterative kmer values from 110 to 150 and from 20 to 150, produced stepwise by 10. Assembly statistics were computed for each metagenome with MetaQUAST v5.0.2

[28]. The N50, number of contigs, and total length of the assemblies were compared for each kmer step, and assembly performance was calculated as the product of the N50 (in kilobases) and read mapping rate (in percent) (Vollmers et al., 2017). The 150-kmer assembly from the 110 to 150 kmer run was chosen as the best assembly using this assembly performance metric.

The 2012 and 2014 metagenomes from Meyer et al. 2016 and Tully et al. 2018 had been previously sheared to 175 bp and sequenced on an Illumina HiSeq 100 at the W.M. Keck sequencing facility at the Marine Biological Laboratory, resulting in an average read length of ~110 bp. These metagenomes were reassembled using the same pipeline as the 2017 metagenomes, but the assemblies were produced for comparison using iterative kmer values from 110 to 150, 90 to 130, and 70 to 100. The 100 kmer assembly step from the 70 to 100 kmer run was chosen as the highest-quality assembly using the assembly performance metric in Vollmers et al. [87] (Supplementary Table 2).

Quality filtering and assembly were performed on the metatranscriptomes in the same manner as for the 2017 metagenomes, and the highest quality assembly (150kmer step) was used for all downstream analyses (Supplementary Table 2).

*Binning and Metagenome Assembled Genomes (MAGs)*

Assembled metagenomes from 2017 and metatranscriptomes from 2012, 2014, and 2017 were prepared for binning with Binsanity [36] by building a bowtie index from each assembly using bowtie2-build version 2.3.4.1 [88]. Sequence alignment map (SAM) files were generated in bowtie from all the North Pond metagenomes (2012-2017) using each bowtie index file. The SAM files were then converted to a compressed binary version (BAM files) using samtools version 1.8 [89]. These BAM files and the assembled metagenomes and metatranscriptomes were run through Binsanity iteratively, a total of six times with a refinement step in between each binning and using CheckM version 1.0.11 [37] to identify high-completion bins. Low-completion and high-redundancy bins were combined after each binning step to be rebinned. Bins were classified using the following parameters:

1) High-completion: >90% complete with <10% redundancy, greater than 80% with <5% redundancy, or >50% with <2% redundancy
2) Low-completion: <50% complete with <5%redundancy
3) Strain heterogeneity: >90% complete with >90% strain heterogeneity
4) High-redundancy: >80% complete with >10% redundancy, or >50% complete >5% redundancy

The FuncSanity function of MetaSanity [44] was used to annotate functional orthologies against the KEGG database [43]; these annotations were run through KEGG-Decoder [90] to determine the completeness of geochemical pathways of interest. Proteases were identified for Prodigal-predict ORFs [91] and searched against the MEROPS database [92] using HMMER v3.2b [93] in MetaSanity, and further analyzed using PSortB v3.0 [94] and SignalP v5.0 [95] to identify signals indicating putative extracellular proteases. Carbohydrate-active enzymes were annotated against the CAZy database [96] within the MetaSanity pipeline.

*Mapping MAGs to Metagenomes and Metatranscriptomes*

A bowtie2 index was built from a concatenated file of all MAG contigs using bowtie2-build. Then, the quality-filtered metagenome reads and metatranscriptome reads were mapped against the MAG bowtie2 index using bowtie2 with the –no-unal flag. The resulting SAM files (containing only the reads which mapped) were converted to BAM files, then BamM (https://github.com/Ecogenomics/BamM) was used to remove all alignments with <95% identity and <75% alignment coverage. The number of reads in each remaining BAM file was counted using Binsanity-profile (v0.3.3) [36]. The read counts were then used to calculate the normalized relative fraction of the metagenomes or metatranscriptomes that mapped to the MAGs using the following equation:

$$\frac{\frac{Reads}{bp}\text{ per genome}}{\sum\frac{Reads}{bp}\text{ all genome}} \times \frac{\sum\text{ Recruited reads to genomes}}{\text{total reads}} \times 100$$

*Discerning Nitrite Oxidation/Nitrate Reduction and Methane/Ammonia Oxidation*

Because genes for nitrite oxidation (*nxrAB*) and nitrate reduction (*narGH*) are annotated under the same KEGG orthology group, we discerned between these processes using three sources of evidence: (1) the taxa annotations of the ORFs called by IMG and mapped by Kallisto (Supplemental Table 4), (2) a phylogenetic tree of *nxrA* and *narG* proteins from the UniProt database [97], and (3) *nxrA* and *narG* protein annotations in the MAGs. A global alignment of *nxrA*/*narG* protein sequences from the MAGs and known *nxrA* and *narG* proteins from UniProt was performed using Geneious v. 9.0.5 (https://www.geneious.com), and a neighbor-joining tree was produced using the Jukes-Cantor genetic distance model and 100 bootstrap replicates (Supplemental Figure 2).

Particulate methane monooxygenase (*pmoA*) was distinguished from ammonia monooxygenase (*amoA*) using the taxa annotations of the ORFs called by IMG and mapped by Kallisto.

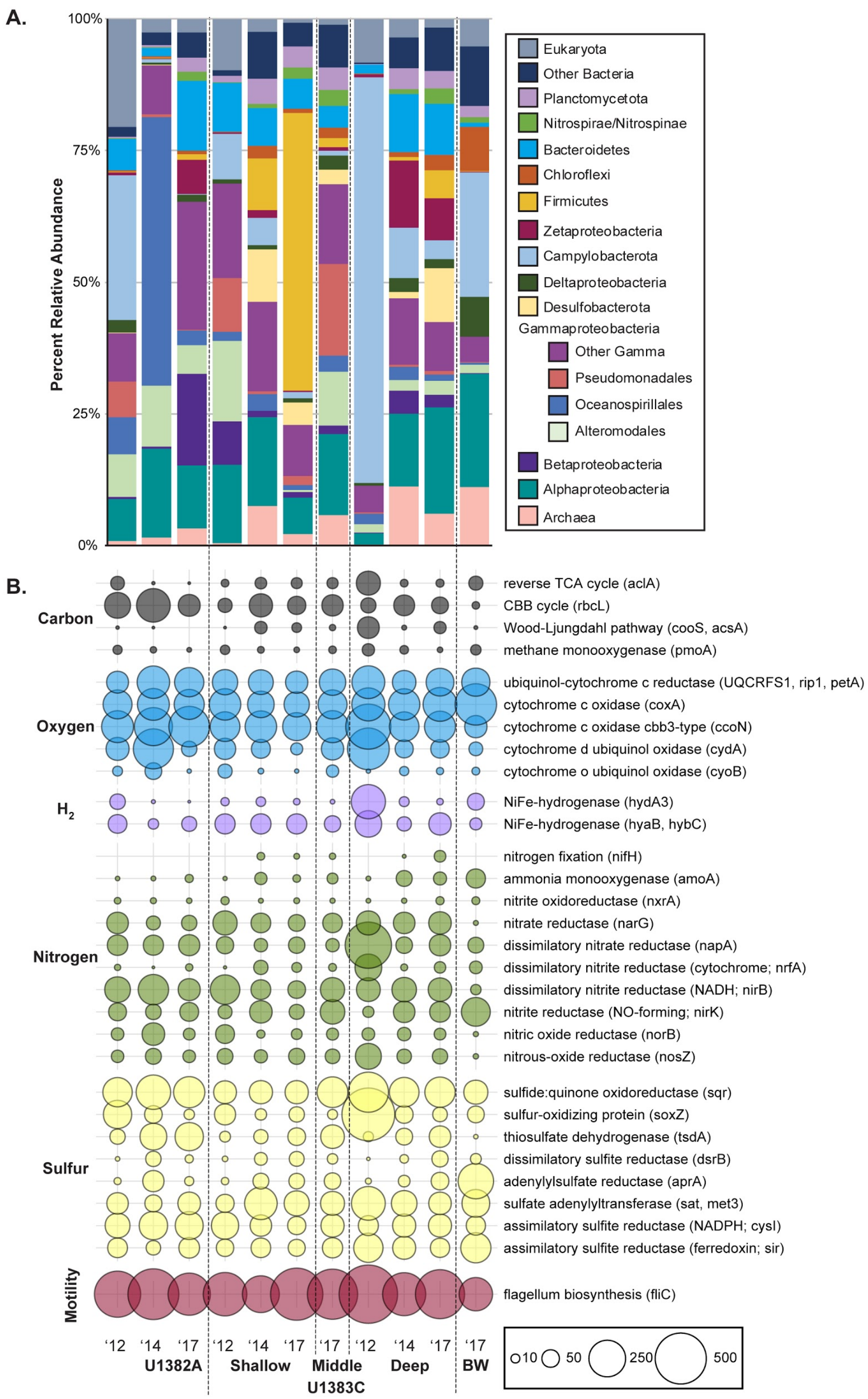


**Supplementary Figure 1.** A) Relative abundance of taxa associated with the small subunit (16S/18S) and large subunit (23S/28S) ribosomal genes annotated in the metagenomes. Ribosomal genes identified using SortMeRNA and annotated using UCLUST. B) Normalized abundance of key genes for carbon, oxygen, hydrogen, nitrogen, sulfur, phosphate, and iron within the metagenome at each site over three sampling years, in transcripts per million reads (TPM).

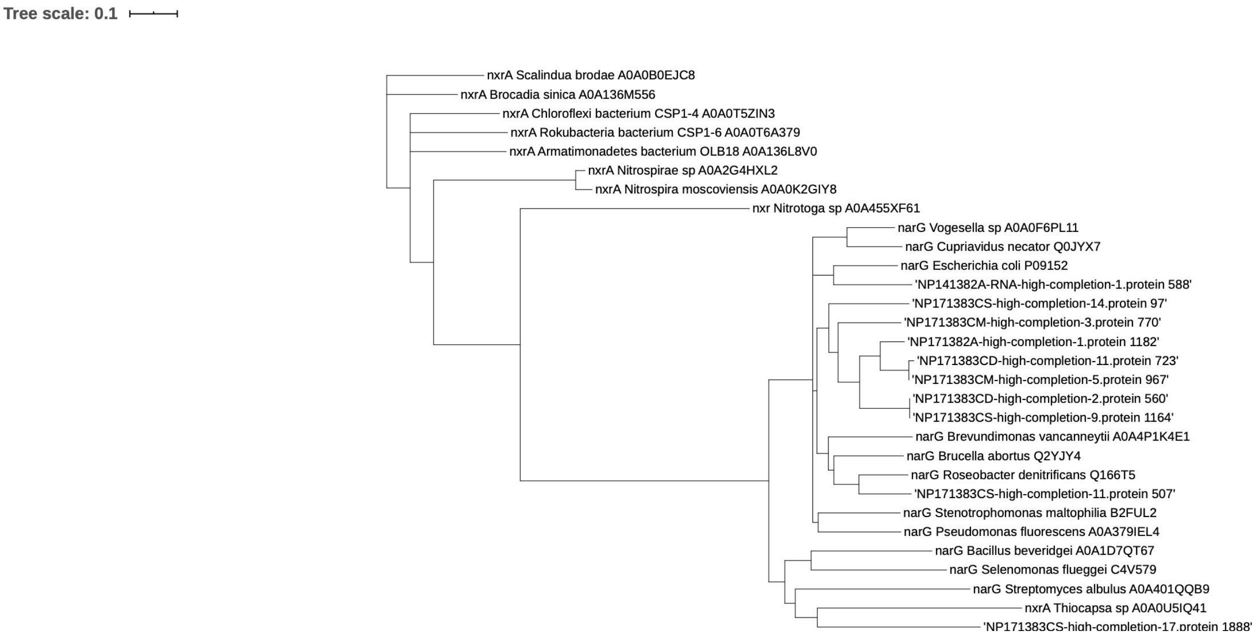


**Supplementary Figure 2.** Bootstrapped phylogenetic tree of nxrA and narG protein sequences annotated in the metagenome assembled genomes (MAGs), including known nxrA and narG protein sequences acquired from UniProt (the Uniprot Consortium, 2019).

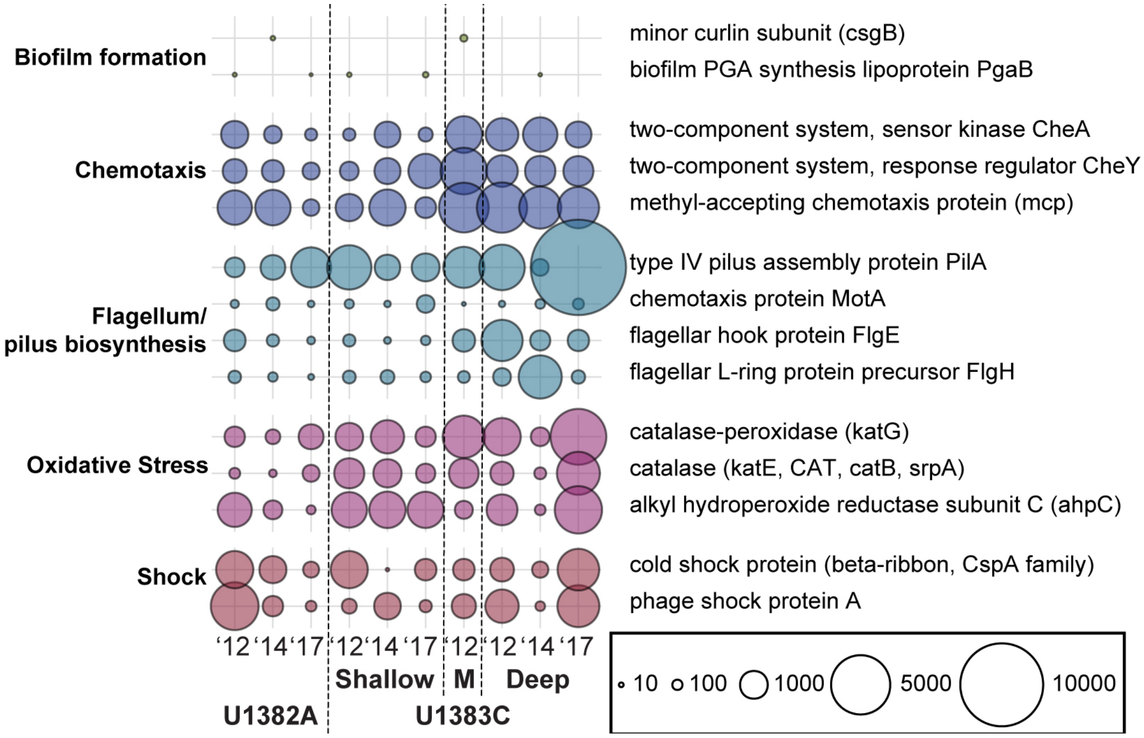


**Supplementary Figure 3.** Normalized transcript abundance of biofilm, chemotaxis, motility, oxidative stress, and shock genes in the metatranscriptomes, in transcripts per million reads (TPM).

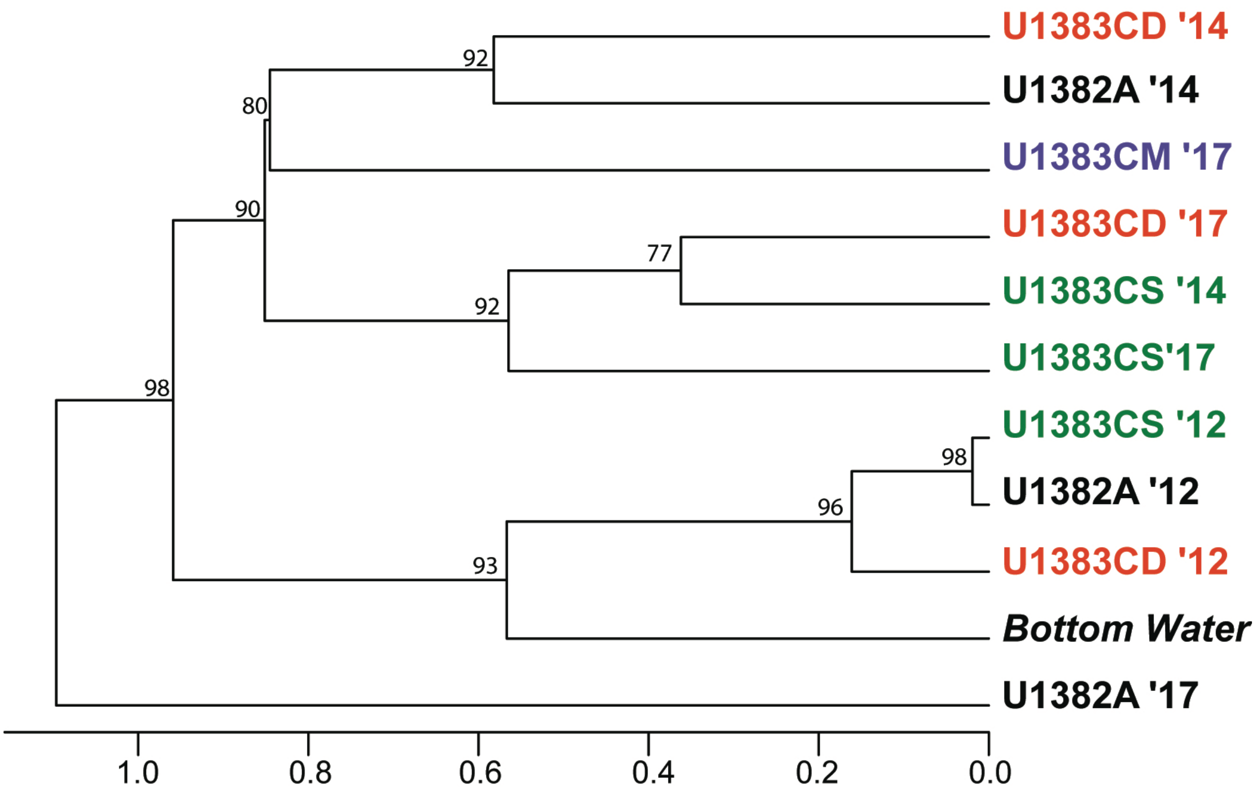


**Supplementary Figure 4.** Hierarchical clustering of MAG abundances in the metagenomes produced using multiscale bootstrap resampling. The scale bar indicates correlation distance between samples.

**Supplementary Tables:**
https://figshare.com/articles/dataset/North_Pond_Metatranscriptomics_Supplemental_Tables/12756248

**Supplementary Table 1.** DNA and RNA extraction yields for all samples (in ng/µl).

**Supplementary Table 2.** Sequence, quality filtering, and assembly data and accession numbers for metagenomes and metatranscriptomes.

**Supplementary Table 3.** Completeness, contamination, strain heterogeneity, and taxonomic identification of the 64 high-completion MAGs obtained by binning.

**Supplementary Table 4.** Counts per million reads (CPM) of all annotated genes in each metatranscriptome.

**Supplementary Table 5.** Completeness of all KEGG modules in all high-completion MAGs determined using Hidden Markov Models with KEGG Decoder. Completeness of each enzymatic pathway is expressed as a percentage (0 to 100%).

**Supplementary Table 6.** Extracellular proteases annotated in all 64 high-completion bins using a Prokka search against the MEROPS database followed by analysis using PSortb and SignalP.

**Supplementary Table 7.** Carbohydrate-active enzymes annotated in all 64 high-completion bins using a Prokka search against the CAZy database.

**Supplementary Table 8.** Genes related to iron acquisition, storage, and reduction/oxidation in each MAG, annotated using FeGenie.
